# Deep learning accurately predicts estrogen receptor status in breast cancer metabolomics data

**DOI:** 10.1101/214254

**Authors:** Fadhl M Alakwaa, Kumardeep Chaudhary, Lana X Garmire

**Author notes:** To whom the correspondence should be addressed: Lana X Garmire, PhD, Associate Professor, Email address.

## Abstract

Metabolomics holds the promise as a new technology to diagnose highly heterogeneous diseases. Conventionally, metabolomics data analysis for diagnosis is done using various statistical and machine learning based classification methods. However, it remains unknown if deep neural network, a class of increasingly popular machine learning methods, is suitable to classify metabolomics data. Here we use a cohort of 271 breast cancer tissues, 204 positive estrogen receptor (ER+) and 67 negative estrogen receptor (ER-), to test the accuracies of autoencoder, a deep learning (DL) framework, as well as six widely used machine learning models, namely Random Forest (RF), Support Vector Machines (SVM), Recursive Partitioning and Regression Trees (RPART), Linear Discriminant Analysis (LDA), Prediction Analysis for Microarrays (PAM), and Generalized Boosted Models (GBM). DL framework has the highest area under the curve (AUC) of 0.93 in classifying ER+/ER-patients, compared to the other six machine learning algorithms. Furthermore, the biological interpretation of the first hidden layer reveals eight commonly enriched significant metabolomics pathways (adjusted P-value<0.05) that cannot be discovered by other machine learning methods. Among them, protein digestion & absorption and ATP-binding cassette (ABC) transporters pathways are also confirmed in integrated analysis between metabolomics and gene expression data in these samples. In summary, deep learning method shows advantages for metabolomics based breast cancer ER status classification, with both the highest prediction accurcy (AUC=0.93) and better revelation of disease biology. We encourage the adoption of autoencoder based deep learning method in the metabolomics research community for classification.

## Introduction

According to Global Health Estimates (WHO 2013), more than half million women died due of breast cancer worldwide^1^. Breast cancer is the second leading cause of cancer-related deaths among women in the United States^2^. Based on human epidermal growth factor receptor 2 (Her2), progesteron receptor (PR) and estrogen receptor (ER), breast cancer can be categorized into four molecular subtypes^3^: Luminal A (ER+, PR+/- and Her2-), Luminal B (ER+, PR+/- and Her2+/-), Her2-enriched (ER-, PR- and Her2+), and triple negative (ER-, PR- and Her2)^4^. The survival outcomes differ significantly among these subtypes. Luminal A and B subtypes have a relatively good prognosis, however triple negative tumors and Her2 tumors have very poor prognosis^5^. Identification of molecular subtypes is crucial in determining cancer prognosis and therapeutic selection. Recently, many studies used metabolomics data to segregate molecular subtypes, given that breast cancer is manifested as a metabolic disease^6, 7^. For example, glutamate-to-glutamine ratio and aerobic glycolysis were proposed as biomarkers of ER and Her2 status, respectively^8, 9^.

Metabolomics studies are usually done by three major platforms: gas chromatography-mass spectrometry (GC-MS), liquid chromatography (LC-MS), and nuclear magnetic resonance (NMR). The parallel use of these instruments allows detecting more metabolites for the same sample. Coupling with the development in the instrumentations, state-of-the-art data analysis tools are much needed to handle the large amount of metabolite data generated. For problems of metabolomics data classification and regression, machine learning algorithms have been applied^10^. For example, Random Forest (RF) is a widely used machine learning algorithm based on decision tree theory. It works with high-dimensional data and can deal with unbalanced and missing values in the data^11^. Support Vector Machine (SVM) is another machine learning algorithm that separates the metabolites data with N data points into (N-1) dimensional hyperplane^12^. SVM was used to classify healthy and pneumonia patients based on nuclear magnetic resonance (NMR) metabolomics data^12^.

DL or deep neural network, is a new class of machine learning methods that have been successfully applied to various areas of genomics research^13, 14^, including predicting the intrinsic molecular subtypes of breast cancer^15^, inferring expression profiles of genes^16^ and predicting the functional activity of genomic sequence^17^. In a recent study, denoising autoencoder (DAs), a type of DL algorithm, was applied to gene expression data of the breast cancer^15^. It successfully extracted features that stratify normal/tumor samples, ER+/ER-status, and intrinsic molecular subtypes. In another study based on gene expression data, DL outperformed linear regression in inference of the expression of target genes from the expression of landmark genes^16^. Moreover, an open source conventional neural networks (CNNs) package “Basset” was developed to learn the functional activity of 164 cell types DNA sequences from genomics data, and to annotate the non-coding genome^17^. Compared to the flourishing applications of DL in genomics, it remains unknown if deep neural network is suitable to classify metabolomics data, esp. when the samples are of medium size (i.e. several hundred).

Here we applied feed-forward networks, a type of DL framework, as an alternative to the machine learning methods such as those listed earlier, to classify metabolomics data. We examined the predictive accuracy of the DL and other machine learning algorithms to predict ER status from a public metabolomics dataset^18^. We demonstrated this DL method performs better than a wide cluster of machine learning methods, including Random Forest (RF), Support Vector Machines (SVM), Recursive Partitioning and Regression Trees (RPART), Linear Discriminant Analysis (LDA), Prediction Analysis for Microarrays (PAM), and Generalized Boosted Models (GBM). Furthermore, the biological interpretation of the hidden layers reveals eight breast cancer related pathways such as central carbon metabolism in cancer and glutathione metabolism. Moreover, we further analyzed the extracted features from our DL model, by mapping the biosynthetic enzymes involved in the metabolomics pathways.

## Materials and Methods

### Data set

The metabolomics data used in this study consists of 271 breast cancer samples (204 ER+ and 67 ER-) collected from a biobank at the Pathology Department of Charité Hospital, Berlin, Germany^18^. Metabolomics profiles of these BC patients can be downloaded from the supporting material of this study^19^. A total of 162 metabolites with known chemical structure were measured using gas chromatography followed by time of flight mass spectroscopy (GC-TOFMS) for all tissues samples. A detailed description of the protocols and the platforms used in this study were described in ^18^. For validation, we downloaded gene expression dataset GSE59198^20^ from the Gene Expression Omnibus (GEO) database, which is composed of 154 samples, a subset of the 271 samples. In this data set, the gene expression profiles of BC tumor tissues (122 ER+ and 32 ER-) were analyzed using the cDNA-mediated Annealing, Selection, Extension and Ligation (DASL) assay. A total of 15,927 genes were detected (p<0.01) in at least 10% of the samples after applying spline normalization. Data can be downloaded from GEO repository <u>http://www.ncbi.nlm.nih.gov/geo</u>.

### Data Preprocessing

We used K-Nearest Neighbors (KNN) method to impute missing metabolomics data^21^. To adjust for the offset between high and low-intensity features, and to reduce the heteroscedasticity, the logged value of each metabolite was centered by its mean 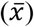 and autoscaled by its standard deviation (*s*) as described in Equation 1^22^. We used quantile normalization to reduce sample-to-sample variation^23^.

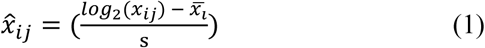

### Deep Learning

DL refers to deep neural network framework, which is widely applied in pattern recognition, image processing, computer vision, and recently in bioinformatics^13, 24, 25^. Similar to other feed-forward artificial neural networks (ANNs), DL employs more than one hidden layer (*y*) that connects the input (*x*) and output layer (*z*) via a weight (*W*) matrix as shown in equation (2). Here we use sigmoid function as the transitioning function.

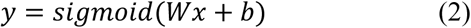

Activation value of the hidden layer (*y*) can be calculated by sigmoid of the multiplication of the input sample *x* with the weight matrix *W* and bias *b*. The transpose of the weight matrix W and the bias *b* can then be used to construct the output (*z*) layer, as described in equation (3).

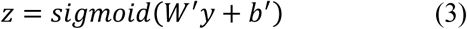

The best set of the weight matrix *W* and bias *b* are expected to minimize the difference between the input layer (*x*) and the output layer (*z*). The objective function is called cross-entropy in equation (4) below, in which the optimal parameters are obtained by stochastic gradient descent searching.

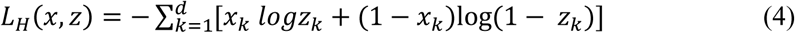

To train the model, we first supplied sample input (*x*) to the first layer and obtained the best parameters (*W,b*) and the activation of the first hidden layer (*y*), and then used y to learn the second layer. We repeated this process in subsequent layers, updating the weights and bias in each epoch. We then used back-propagation to tune the parameters of all layers. Finally, we fed the output of the last hidden layer to a softmax classifier which assigned new labels to the samples^26^. We used *h2o* R package to tune the parameters of the DL model^27^.

### Other machine learning algorithms

We selected a representative set of six machine-learning algorithms that are highly recommended by the metabolomics community and applied widely in the literature reports: Random Forest (RF), Support Vector Machines (SVM), Recursive Partitioning and Regression Trees (RPART), Linear Discriminant Analysis (LDA), Prediction Analysis for Microarrays (PAM), and Generalized Boosted Models (GBM). To get the optimal predictions, we used the *caret* R package^28^ to tune the parameters in the models.

### Modeling and evaluation

We randomly split metabolomics samples into 80% training set and 20% testing set. The 80/20 split is a common practice of splitting ratio for samples of a moderate size in the machine learning applications. We chose this ratio in order to having enough training samples to build a good model and sufficient testing samples to evaluate the model. We performed 10-fold cross-validation on the 80% training data during the model construction process, and tested the model on the hold out 20% of data. We used pROC R package^29^ to compute area under the curve (AUC) of a receiver-operating characteristic (ROC) curve to assess the overall performance of the models. To avoid sampling bias, we repeated the above splitting process ten times and calculated the average AUC on the hold out 10 test samples. To control overfitting, we used two regularization parameters: L1, which increases model stability and causes many weights to become 0 and L2, which prevents weights enlargement.

We tuned DL model and other machine learning algorithms, on the following parameters: DL model: Epochs (number of passes of the full training set), *l*1 (penalty to converge many weights to 0) and *l*2 (penalty to prevent weights enlargement), and input dropout ratio (ratio of ignored neurons in the input layer during training), number of hidden layers; RPART model: complexity parameters (cost of adding node to the tree); GBM model: number of trees and interaction depths; SVM model: cost of classification; RF model: number of trees to fit; PAM model: threshold amount by for each of the class's centroid shrinking towards the all classes’ centroid.

### Feature importance

Features importance was estimated based on model based approach^28^. In other words, a feature is considered important if it contributes to the model performance^30^. We used the variable importance functions varimp in *h2o* and varImp in *caret* R packages, to evaluate the top 20 features.

### Identifiers standardization and differentially expressed genes

We used the PubChem Identifier exchange service^31^ to convert metabolites into their corresponding KEGG compound IDs; we then used KEGG API^32^ to get the compound pathways and enzyme IDs. We used *limma* R package^33^ to find enzymes with high fold changes as well as significant adjusted p-values between ER+ and ER-samples.

### Metabolomics enzymes network reconstruction and visualization

We used MetaScape^34^ v3.1.3, a Cytoscape plug-in to generate gene-metabolite network which integrates reaction and pathway information from KEGG and Edinburgh human metabolic network (EHMN) databases. To build enzyme-metabolite network, we selected a pathway based network from Metsacpe analysis options. The input of this step were two files. The first file included the compound KEGG IDs, p-value and the fold change values of the top 20 metabolites extracted from the DL model. The second file included the enzyme KEGG IDs, p-value and the fold change values of the 898 genes whose expression values were statistically significantly different between ER- and ER+ samples.

### Metabolites enzymes correlation

We calculated the correlations between the intensity levels of the metabolites and enzymes using Spearman's Correlation Coefficient in R. We plot the Circos plot of the strongest correlation using Circlize R package v0.4.0.

### Joint significant pathway analysis

To perform joint significant pathway analysis on metabolomics and gene expression data from the same samples, we considered a comprehensive list of pathways from Reactome, EHMN, and KEGG databases, using online web tool IMPaLA35, and calculated hypergeometric p-values of genes (*P*_*G*_) and metabolites (*P*_*M*_). The joint P-value (*P*_*j*_) between metabolites and genes for pathway *i* was calculated as *P*_*ji*_ =*P*_*Gi*_ *P*_*Mi*_ ^36^. This value was adjusted to control for multiple testing with the False Discovery Rate method.

### Code availability

We include all preprocessing and the learning steps of the DL method as an R script in the supplementary file 1.

## Results

### Workflow of autoencoder based classification

We aim to assess the predictive ability of the DL framework to separate breast cancer patients based on their ER status, using metabolomics data. Towards this goal, we implemented the workflow of DL framework as in **Figure 1**. We applied preprocessing steps (log transformation, centering, autoscaling, and quantile normalization) before constructing the DL model, as recommended by others^18, 22^. Before training the model, we pre-trained the model using autoencoder and the whole data without labels. This step improves the model performance, avoids random initialization of the weights, and selects the best model architecture^37^. Then we trained the DL model using a wide range of parameters and selected the best model with the minimum mean square error (see Materials and Methods).

**Figure 1:**
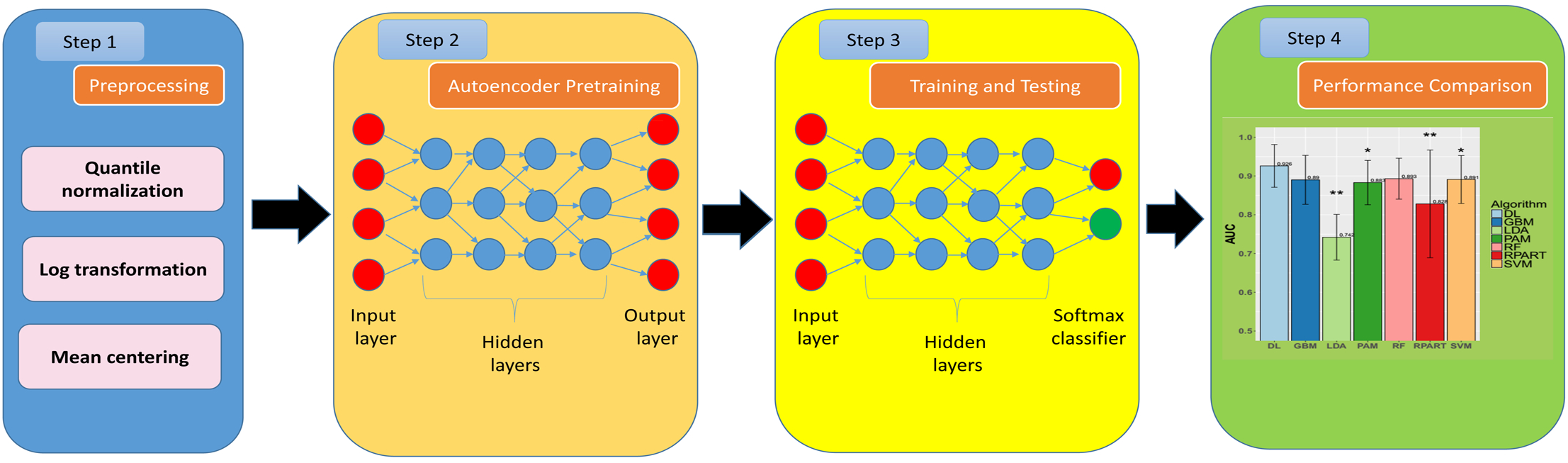
Block diagram of the proposed system. The first step is the preprocessing (log transformation, centering, autoscaling and quantile normalization). We used Autoencoder pretraining (unsupervised step) to initial model weights and select model architecture. Model used the 80% of data split to train the model and the remaining 20% to measure model performance. The data was split 10 times to avoid the bias of data sampling, and the average AUC was calculated on the 10 holds out test samples.

### Performance of the autoencoder based deep learning classification

We compared DL with six other machine-learning methods commonly used in the community: Random Forest (RF), Support Vector Machines (SVM), Recursive Partitioning and Regression Trees (RPART), Linear Discriminant Analysis (LDA), Prediction Analysis for Microarrays (PAM), and Generalized Boosted Models (GBM). To assess the predictive power of the models, we partitioned the data into 80% training and 20% testing subsets. We performed 10-fold cross-validation on the 80% training data, and tested the model on the hold out 20% of data. To avoid sampling bias, we performed 10 independent splitting of training and testing subsets. We reported the averaged AUCs calculated on the hold out test sets. As shown in **Figure 2A**, the average AUC of DL yields the best AUC of 0.93, compared to other six classification methods. The superiority of DL accuracy is statistically significant (Wilcoxon signed-rank test P<0.05) than other methods, except RF and GBM. LDA and RPAT had the worst accuracy, likely due to their sensitivity to overfitting and unfit to the non-linear problems^38^.

**Figure 2:**
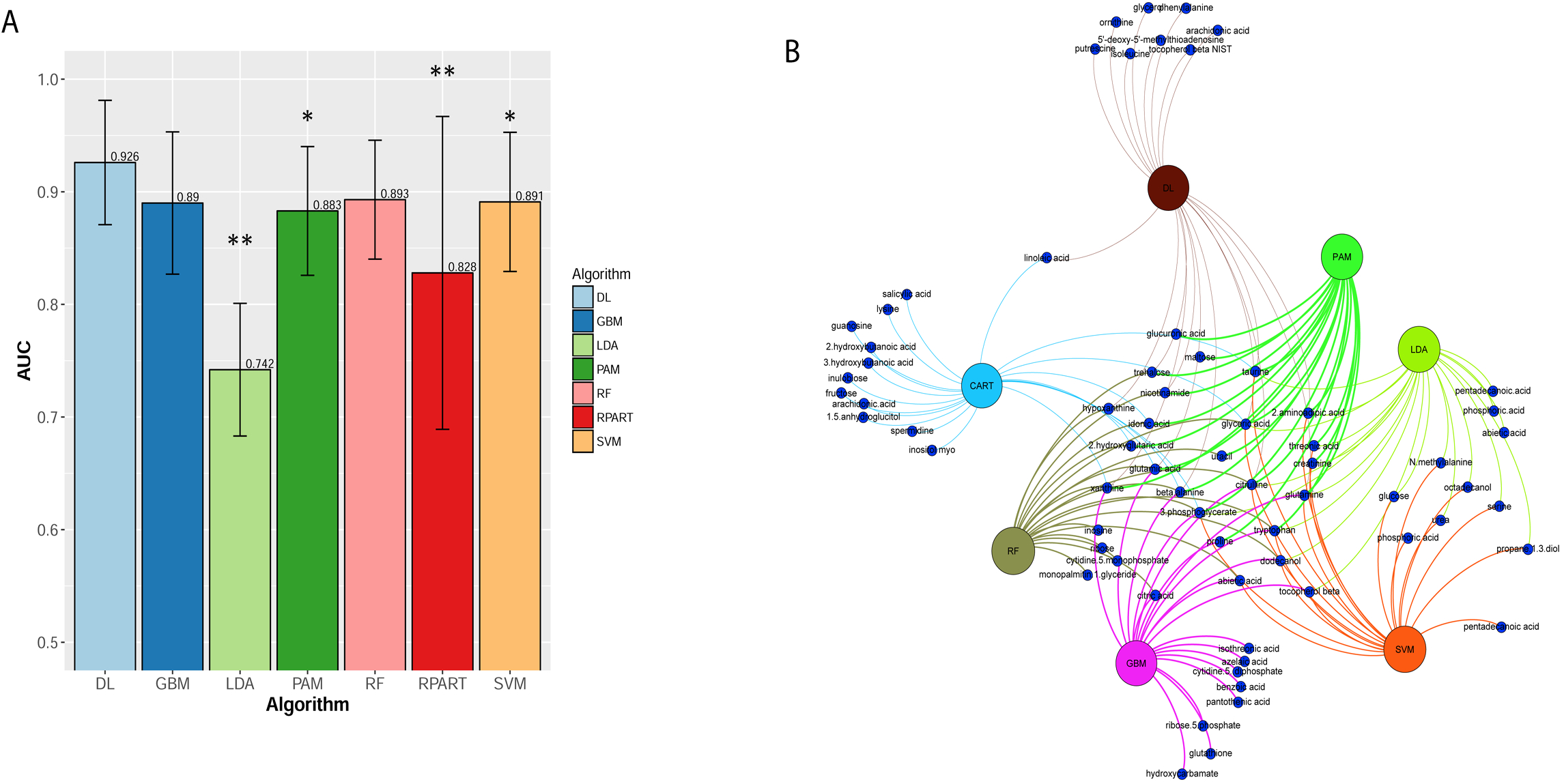
**A:** The average AUC on 10 hold out test samples of the DL framework against six machine learning algorithms for prediction of ER status from metabolomics data: Recursive Partitioning and Regression Trees (RPART) (0.83), Linear Discriminant Analysis (LDA) (0.74), Support Vector Machine (SVM)(0.89), DeepLearning (DL)(0.93), Random Forest (RF)(0.89), Generalized Boosted Models (GBM)(0.89), and Prediction Analysis for Microarrays (PAM)(0.88). The above algorithms were run 10 times on different train/test splits. We used pairwise Wilcoxon signed-rank test to estimate the statistical significance of the difference in performance between DL and other methods (** p<0.01, * p<0.1). **B:** Bipartite graph of the top 20 important metabolites extracted from DL model and other machine learning algorithms. Large nodes represent the models and small nodes are metabolites. A connection between metabolite and the model means this metabolite is one of the top 20 high importance metabolites extracted by this model.

DL as other machine learning algorithm needs more samples to achieve high accuracy^39^. To assess the effect of sample size on various models, we randomly removed ¼, ½, and ¾ of the data sets (**Figure S1**). As expected, decreasing in sample size decreases the averaged AUCs of all classification methods in general except LDA on ¼ samples, due to overfitting. Notably, the reduction of average AUC in DL is most pronounced among all methods, from the full to ¾ data set (**Figure S1**). While DL loses the best average AUC status when the sample size is around 255, GBM, SVM and RF have the highest AUC for small sample sizes of 203, 136 and 68, respectively. Similarly, we also experimented the effect of metabolite size on various models (**Figure S2**). We randomly removed ⅛, ¼, and ½ of the 162 metabolites. Even with reduced numbers of metabolites, deep learning and the robust machine learning method SVM still have fairly good predictions, compared to other algorithms tested. This suggests that, due to colinearality, much of information still exist in the remaining metabolites. Together, the drop-out experiments (**Figures S1 and S2**) demonstrate that DL method is sensitive to sample size, but much less sensitive to metabolite size.

### Important features from DL

To relate the importance of metabolites to ER status directly, we ranked the metabolites extracted from DL model based on their functional contributions to the outputs. In this approach, features that provide unique information to the trained network are ranked more importantly than those giving redundant information^40^. We listed the top 20 metabolites from DL in Table S1, and presented their heatmap and boxplots in **Figure S3**. Note the choice of 20 metabolite is guided by the original study, in which 19 out of 162 metabolites were claimed to change significantly among training and validation samples^19^. The original author divided the 271 samples into two parts, the training (2/3) and the validation (1/3) set. Among the training set, 65 metabolites were different in ER- and ER+ and only 19 metabolites were validated in the validation set.

Among the 20 features, the top five features are beta-alanine, xanthine, isoleucine, glutamate, and taurine. These five metabolites have been either proposed as breast cancer biomarkers or associated with breast cancers in the original metabolomics report^19^ and/or other studies^6, 8, 41–43^. For instance, Budczies et al. ^19^ found that beta-alanine had the most significant and largest fold changes between ER-(n=67) and ER+ (n=204) tumor tissues. In another study, Glutamate was suggested as markers to segregate ER- from ER+ in the training (n=186) as well as validation dataset (n=88)^8^. Glutamate to glutamine ratio (GGR) was significantly increased in the ER-tumors as compared to ER+. Overall survival analyses suggested GGR as a positive prognostic marker for BC^8^. In another study, Fan et al. classified BC plasma samples into subtypes i.e. ER+ vs ER- and HER2+ vs HER2-, based on a training set (n=51) and another test set (n=45)^6^. They found isoleucine had significant differential level between ER+ (lower) and ER-(higher) samples. Similarly, a study among female breast cancer patients (n=50) suggested serum taurine as an early marker, where its level was significantly lower than the normal (n=20) and high risk samples (n=15)^42^. In a cell line based study, xanthine was suggested as potential biomarker of breast cancer metastasis^43^, as it had the highest variable influence on projection (VIP) in the three pair-wise comparisons among MCF-7/MCF-10A, MDA-MB-231/MCF-10A and MDA-MB-231/MCF-7^43^.

Further, we compared DL top 20 features with the same number of top features from all other methods in a bipartite graph (**Figure 2B**). Twelve metabolites are shared between DL and one or more algorithms. Among them, 1 (xanthine) is shared by six methods, 2 (glyceric acid and citrulline) are shared by five methods, 4 (glutamine, taurine, glutamine acid, and beta-alanine) are shared by four methods, 1 (2-aminoadipic acid) is shared by three methods, 2 (nicotinamide acid and trehalose) are shared by two methods, and two (linoleic acid and hypoxanthine) are shared by one method (Table S1). Additionally, DL has identified 8 unique metabolites: isoleucine, putrescine, glycerol, 5'-deoxy-5'-methylthioadenosine, ornithine, tocopherol beta, phenylalanine, and arachidonic acid,

### The biological relevance of the hidden layers

To understand the high performance of the DL model, we probed into the hidden layer and analyzed the 25 activation nodes from the first hidden layer. Among the top 12 nodes with the variances > 0.1, node 8, 22 and 25 are significantly correlated with the samples’ ER-status (P=1.14e-12), whereas all other top 9 nodes are associated with the ER+ status (**Figure 3A**). These results confirm that the nodes in DL have significant biological meaning.

**Figure 3:**
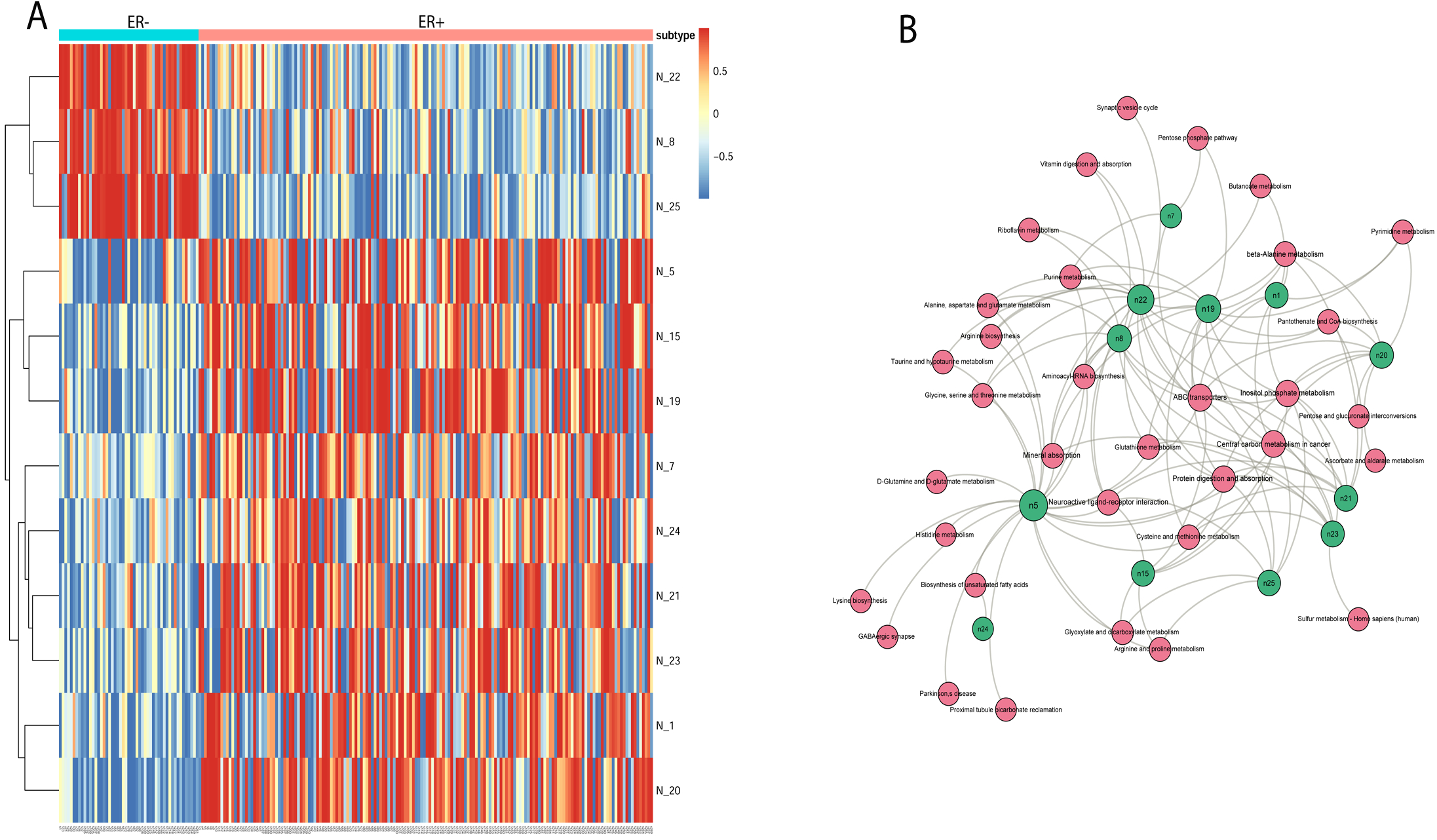
Biological relevance of the DL hidden layers. (A) Activation levels of the high variance nodes extracted from the layer 1 of the DL model. Columns are samples and rows are the top 12 nodes with high variance > 0.5. (B) Bipartite graph of enriched significant metabolomics pathways and top hidden nodes. The nodes represent enriched pathways common to all top 12 nodes (green color) in the 1^st^ hidden layer of DL in KEGG pathway enrichment analysis (FDR< 0.05).

We identified a total of 129 metabolites which contribute most to the activation values of the top 12 nodes. Their relationships between the 129 metabolites and 12 nodes are shown in **Figure S4**. We define that metabolite *x* contributes to the activation value (*y*) of node *n*, if the aboslute value of the weight connecting metabolite *x* and node *n* is greater that 0.1. Beta-alanine and xanthine are the most common metabolites from all top 12 nodes. Among nodes 8, 22, and 25 which are highly correlated with ER-(**Figure 3A**), four common metabolites are shared: inositol, glutamate, xanthine, and uracil. Xanthine was among the panel of prognostic markers of breast cancer metastasis based on the metabolic profiling of the three breast cancer cell lines^43^. Glutamate have been reported as biomarkers to segregate ER-from ER+ in the training as well as validation dataset, as described earlier^8^. Inositol phosphate metabolism pathway was previously reported to be associated with breast cancer, but not between ER+ and ER-cancers^44^. Uracil is, however, a potencial new marker for ER-breast cancer that was not reported previously, according to our knowledge.

To link the metabolites in **Figure S4** with biological functions, we conducted pathways enrichment analysis using online web tool IMPALA^35^. The pathways are taken from Reactome, EHMN, and KEGG databases. Eight significant breast cancer related pathways (Figure 3B) are enriched in all nodes: protein digestion and absorption, central carbon metabolism in cancer, neuroactive ligand receptor interaction, ABC transporters, mineral absorption, inositol phosphate metabolism, glutathione metabolism, and cysteine and methionine metabolism. Albeit the name of “Neuroactive ligand-receptor interaction”, this pathway is significantly enriched (q-value=0.001) and it was shown changed in breast cancer cell lines ^45^ and naked mole rat ^46^. Aspartate, glucine, taurine and glutamate are metabolites associated with this pathway in the metabolic dataset. Another interesting pathway with the name “mineral absorption” also shows significance (q-value=7.51E-06), attributed by five metabolites tryptophan, alanine, glycine, phosphoric acid, glutamine. All these five metabolites were found related with breast cancer previously^47–49^.

### Integration of DL metabolites and enzymes

We further aimed to validate the important metabolite features of DL model, by integrating metabolomics and gene expression data from the same patients. Towards this, we first conducted a joint pathway analysis between 20 metabolites extracted from DL model and 898 significantly differentiated enzymes between ER+ and ER-samples, using IMPALA (Figure 4). Most of the top significant pathways are related to metabolism of amino acids or protein digestion and absorption. Two pathways remain significant in joint pathway analysis, by comparing to metabolomics based pathway analysis in Figure 3B: protein digestion & absorption and ABC transporters, with 6 and 9 metabolites over-represented respectively. Specifically, urea, inositol allo-, phosphoric acid, glucose, glutamine, Isoleucine, and glutathione are the associated metabolites in ABC transporters. For protein digestion, glutamine, lysine, isoleucine, and beta-alanine are associated metabolites. Some literature evidence shows that protein digestion and ABC transporters are related to breast cancer. For example, humans have 49 members of the ATP-binding cassette (ABC) membrane proteins^50^. Several of them such as ABCB1 and ABCC1 have developed a resistance to drug “multidrug resistance” (MDR) in breast cancer, when they are over-expressed over a period of time^51^.

**Figure 4:**
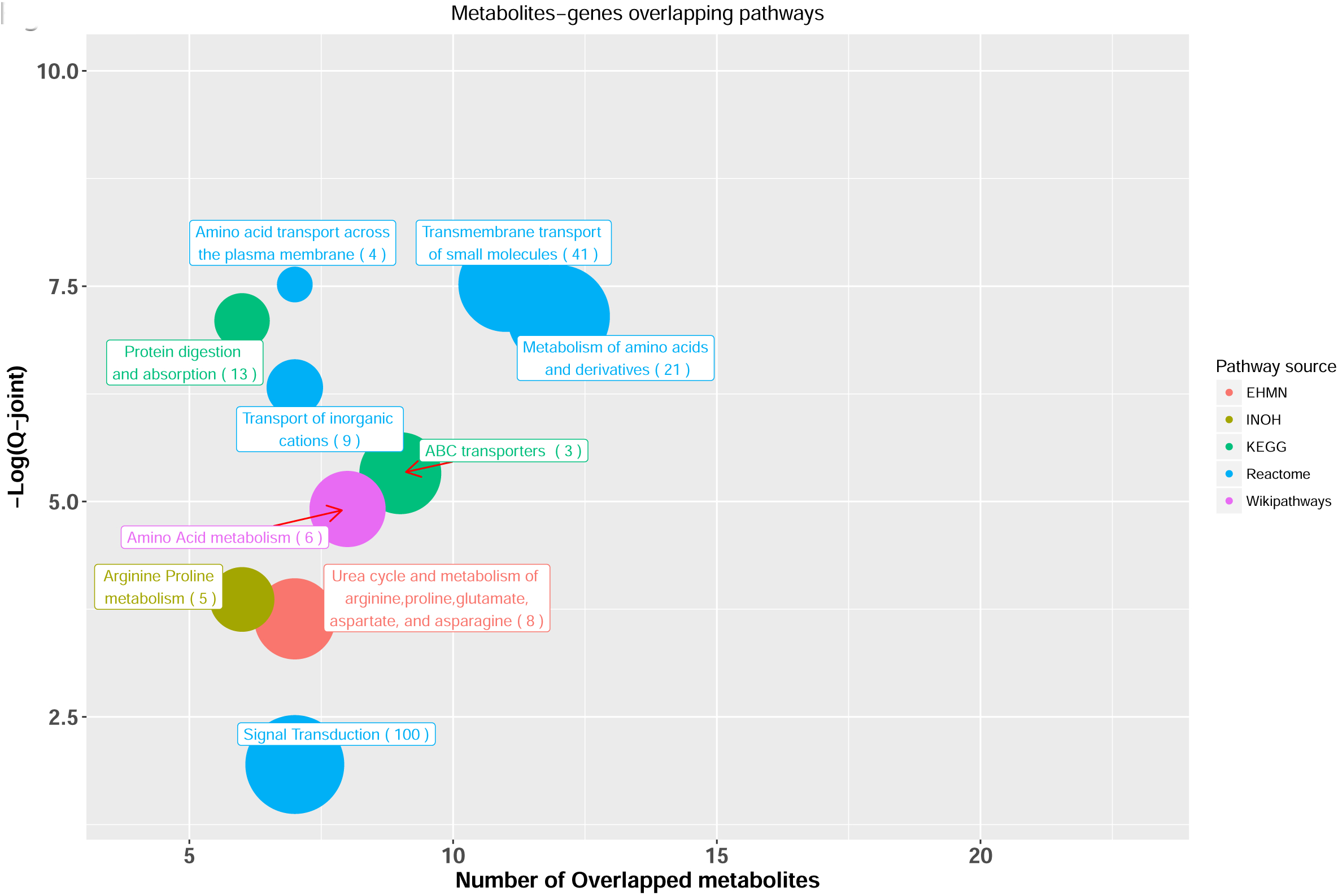
The joint pathway analysis between the top 20 DL metabolites and the high differentiated enzymes. Only significant pathways with at least 5 overlapping metabolites are shown. X-axis shows the number of overlapped metabolites with the number of genes (number in parentheses) involved in the same pathway, y axis shows the adjusted joint *P*-value calculated from IMPALA tool^42^. The size of the nodes represents the size of metabolomic pathway (number of metabolites involved in that pathway). The color of the nodes represents the database source of these pathways.

To gain insights at individual metabolite/enzyme level, we then calculated Spearman correlations between the intensity levels of the top 20 metabolites and enzymes whose gene expression levels are significantly different between ER+/ER-for the same patients^20^. The Circos plot in **Figure 5** shows the names of metabolomics and enzymes that have correlations (|r| > 0.35). Impressively, beta-alanine, the top ranked metabolite in DL, is the single most connected metabolite, correlated to more than 100 significantly differentially expressed enzymes. Pathway analysis of these enzymes correlated with beta-alanine shows strikingly significant enrichment (adjusted p-value =3.84e-05) with FOXM1 transcription factor network pathway. FOXM1 is highly expressed in ER-samples and with a correlation coefficient r=0.5 with beta-alanine.

**Figure 5:**
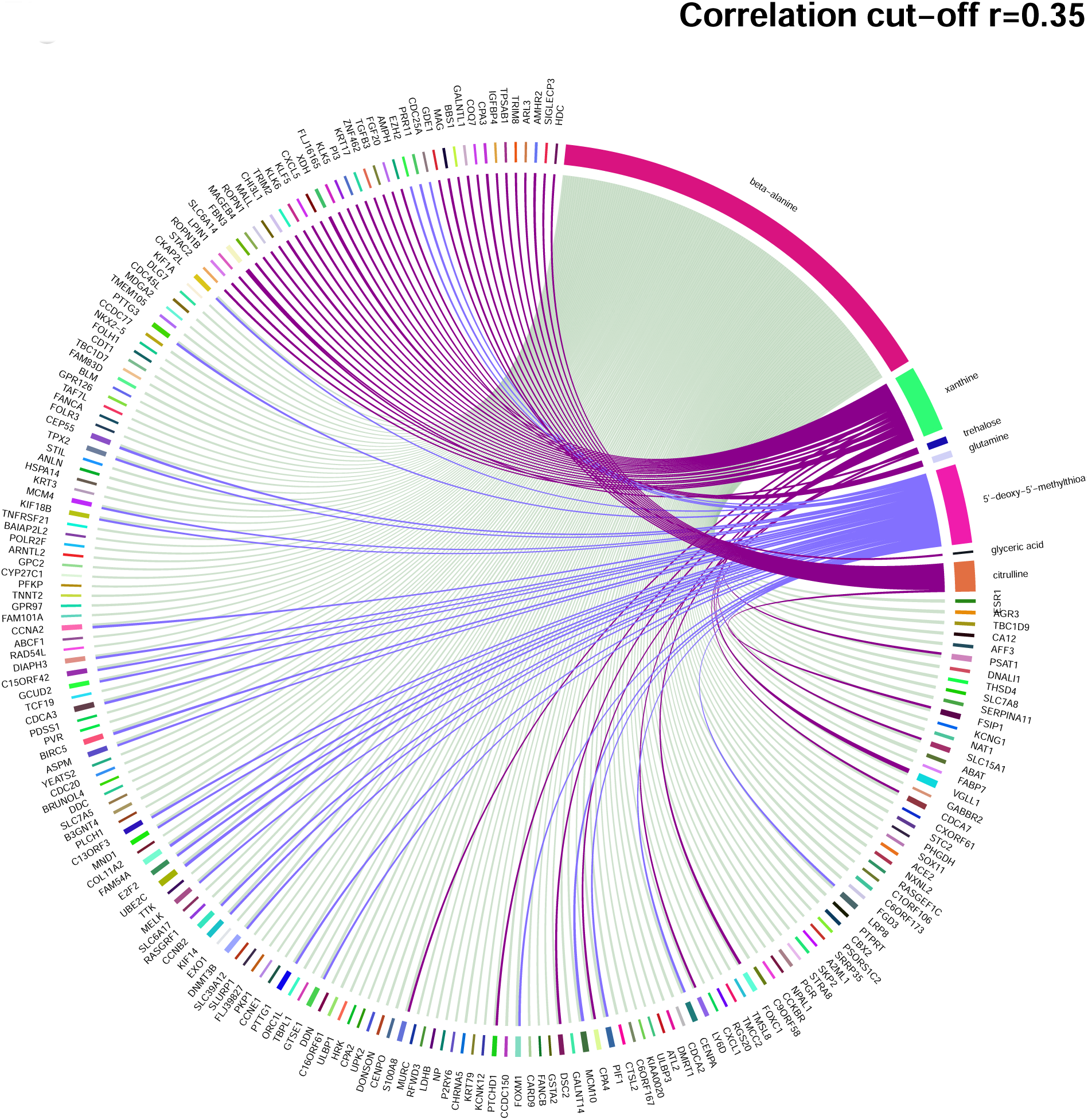
Circos plot of Spearman correlation values between 20 top DL metabolites and high differentiated enzymes with cut-off=|0.35|.

Complementary to the correlation based analysis, we also used Metscape (Cytoscape plug-in) for gene-metabolite network analysis, by combining the ER+/ER-metabolomics data^18^ and gene expression (from GSE59198)^20^ for the same patients. ABAT, the enzyme that catalyze beta-alanine to malonate semialdehyde (Figure 6B), is highly correlated with beta-alanine (r=-0.62, Figure 6A). To understand better the connection between beta-alanine and FOX genes family, we performed motif enrichment analysis for the enzymes interacted with beta-alanine in Figure 6B using PASTAA tool^52^. Interestingly, FOXO1 was one of most significant transcription factors (p= 5.89e-04) that targeted the promoters regions of beta-alanine interacted enzymes.

**Figure 6:**
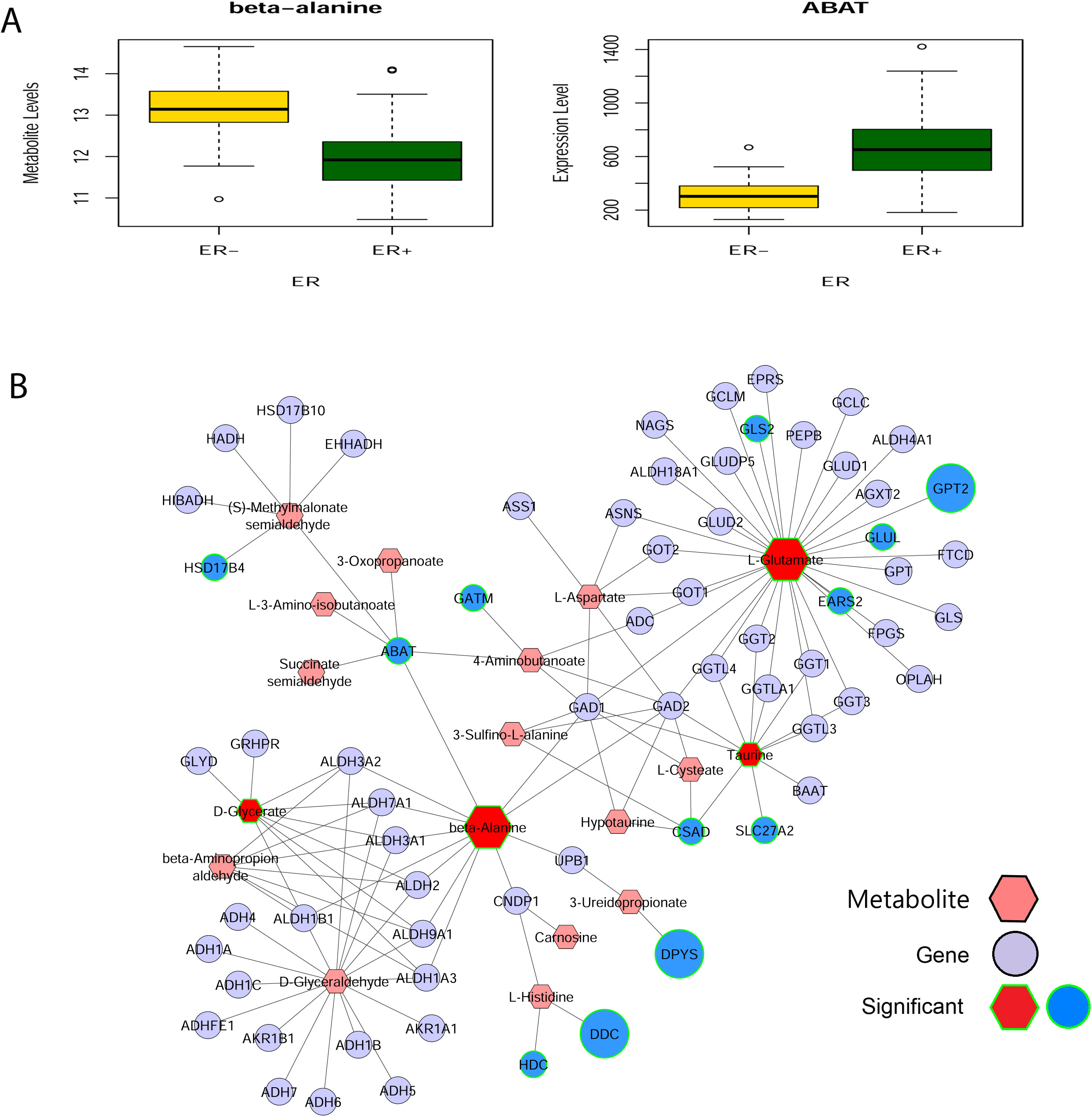
Beta-alanine and ABAT interaction network. (A) Metabolite level of beta-alanine and expression of ABAT. (B) Beta-alanine-ABAT interaction network in ER– breast cancer tissues compared to ER+ breast cancer tissues. Metscape, a Cytoscape plug-in, was used to integrate ER+/ER-metabolomics and gene expression data (GSE59198) of the same patients. Fold change of metabolites (hexagon nodes) or enzymes (circle nodes) are represented by the size of the nodes. The input of Metascape are the top 20 metabolites from the DL model and the 898 genes whose expression values are statistically significantly different between ER- and ER+ samples. Enzymes and metabolites of significant difference are marked by green line(s) on the shapes.

## Discussion

Metabolomics has become a new platform for biomarker discovery. Accompanying this technology, robust and accurate classification methods to predict sample labels are in critical need. Recently, DL methods have gained much attention in domains such as genomics and imaging analysis. However, there has not been any systematic investigation of DL methods in the metabolomics space. In this report, we aimed to fill this void and assessed the performance of feed-forward network, a widely used DL framework, on classifying ER+/ER-breast cancer metabolomics data.

There are many advantages of DL over shallow machine learning algorithms, which are beyond the scope of this study. The conventional machine learning algorithms require engineering domain knowledge to create features from raw data, whereas DL automatically extracts simple features from the input data using general purpose learning procedure. These simple features are mapped into outputs using a complex architecture composed of a series of non-linear functions “hierarchical representations,” to maximize the predictive accuracy of the model optimally. By increasing number of layers and neurons per layers, robust features may be constructed, and error signals can be diminished as they pass through multiple layers^13^. Therefore, DL succeeds to construct high-level transformed features from input data, making it more desirable than shallow machine learning algorithms in this respect^14^.

We demonstrated that DL has a higher predictive accuracy over the other six popular machine learning methods, in detecting ER status from metabolomics data. DL exploits the idea that the higher “succeeding” layer is learned from the lower “preceding” layer and selects the essential metabolites from DL model. These metabolites are useful for the learning process and explain the high predictability of DL compared to conventional machine learning algorithms. DL extracted features that could be considered as novel biomarkers, such as uracil, which were not previously reported as breast cancer. Also, unlike other machine learning methods, DL method offers additional insights on eight KEGG pathway being significantly different due to ER status. All these new observations warrant further investigation.

An interesting new link we discover lies between FOXM1 family and beta-alanine. A recent study showed FOXM1 to be a major cause for resistance to various chemotherapeutics^53^, and reduction of FOXM1 levels induced apoptosis of breast cancer cells^54^. The motif enrichment analysis of the beta-alanine interacted enzymes indicates that the transcription factor FOXO1 targeted the promoter regions of these enzymes. Thus the relationships among beta-alanine, FOXM1 and FOXO1 is worth further investigation. In addition, we found many interesting involvement of DL unique metabolites in breast cancer diagnosis and treatment.

For example, phenylalanine is found significantly elevated in the advanced metastatic breast cancer^55^ and linoleic acid has been used to lower the risk of breast cancer^56^. Also, Putrescine has been known to play a critical role in many metabolomics processes in breast cancer, such as apoptosis, and proliferation^57^. The knock-down experiments on ornithine decarboxylase (ODC), an enzyme which converts ornithine to putrescin, showed the growth inhibition in the ERα+ MCF7 and T47D and ERα-MDA-MB-231 breast cancer cells^58^. Arachidonic acid was previously shown to be integral part of the new signaling for the cell migrations in the MDA-MB-231 breast cancer cells^59^.

Despite the outstanding performance of DL methods, one should be mindful of several caveats in its application in metabolomics research. DL is time-consuming computation (Table S2), relative to some other machine learning methods^40^. Also, metabolomics data sets are generally small, in comparison to imaging data. Thus very small data sets may not be suitable for DL. We experimented with the effects of reducing sample size and metabolite size on the seven methods in comparison, and found that DL is indeed sensitive to the sample size of the study. On the contrary, due to colinearality among metabolites, autoencoder has fairly robust predictions even when the number of metabolites are reduced. Another point of consideration is the reproducibility of the technology itself. A platform with better reproducibility is expected to yield biomarker models that predict more accurately in validation datasets (less overfitting). We thus speculate that DL models based on NMR metabolomics data (more metabolites and better reproducibility) will be more accurate than DL models based on LC-MS data, when other conditions are the same.

Lastly, in this report we compared the ML vs DL under the topic of classification of metabolomics data. The advantages of DL on other non-classification problems in metabolomics research are yet to be explored. For example, unsupervised machine learning algorithms such as PCA and hierarchical clustering were applied to the metabolomics data^60^, and our group is currently exploring using autoencoders for unsupervised learning in metabolomics data. As another example, we have also worked on prognosis prediction using shallow and deep neural network models in the genomics space ^61, 62^. We successfully used autoencoder to integrate multiple omics datasets (RNA-Seq, microRNA-Seq and DNA methylation) to predict patient survival robustly, exemplified by liver cancer [2]. Compared to genomics data, metabolomics data have higher multicolinearity and noise levels. Also the number of identifiable metabolites are lower than the identifiable genes in genomics assays. These issues pose potential challenges when extending genomics tools for metabolomics research. Nevertheless, it will be very interesting to test these DL and neural network models on appropriate metabolomics data sets alone, or in combination with coupled genomics data.

## Conclusions

We show evidence that DL outperforms other machine learning algorithms for ER status classification in breast cancer metabolomics data. The biological interpretation of the hidden layer of the DL model also reveals eight significant breast cancer related pathways, which are not able to obtain from the other machine learning algorithms in comparison.

## Author Contributions

LXG and FMA envisioned the project and designed the work. FMA coded the project and conducted the analysis. KC mapped metabolites and enzymes into KEGG pathway. FMA wrote the manuscript with help from LXG and KC. LXG, FMA and KC have read, revised and approved the final manuscript.

## Acknowledgements

This research was supported by grants K01ES025434 awarded by NIEHS through funds provided by the trans-NIH Big Data to Knowledge (BD2K) initiative (http://datascience.nih.gov/bd2k), P20 COBRE GM103457 awarded by NIH/NIGMS, and R01 LM012373 awarded by NLM, R01 HD084633 awarded by NICHD to LX Garmire. We thank all members in Garmire group for reviewing and commenting the manuscript.

## Supplementary Materials

**Figure S1:** (A) The effect of sample size on the performance of the DL and other machine learning algorithms.

**Figure S2:** The effect of metabolite size on the performance of the DL and other machine learning algorithms.

**Figure S3:** DL 20 top important metabolites. **A.** Heatmap and **B.** Box plot of the 20 top important metabolites extracted from the DL model.

**Figure S4:** Heatmap of the metabolites (columns) which most contribute to the activation value of the top hidden nodes (rows).

**Table S1:** The list of the top 20 important features

**Table S2:** Running time of the seven algorithms on the metabolomics dataset

**Supplementary file 1**: R code of the preprocessing, models training and testing

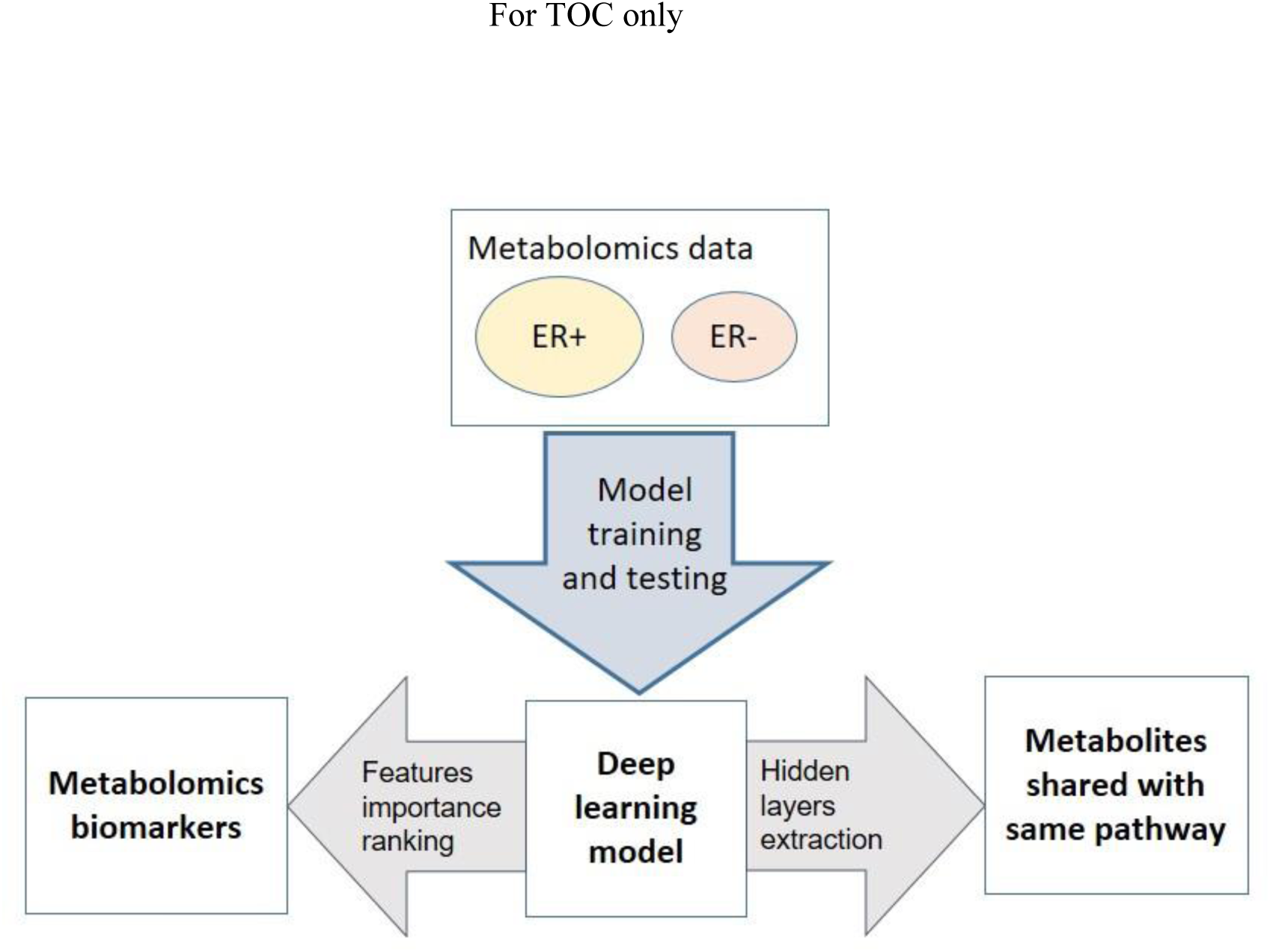

